# The antimicrobial peptide DGL13K is active against resistant gram-negative bacteria and subinhibitory concentrations stimulate bacterial growth without causing resistance

**DOI:** 10.1101/2020.05.08.085282

**Authors:** Sven-Ulrik Gorr, Hunter V. Brigman, Jadyn C. Anderson, Elizabeth B. Hirsch

## Abstract

Antimicrobial peptides may be alternatives to traditional antibiotics with reduced bacterial resistance. The antimicrobial peptide GL13K was derived from the salivary protein BPIFA2. This study determined the relative activity of the L-and D-enantiomers of GL13K to wild-type and drug-resistant strains of three gram-negative species and against *Pseudomonas aeruginosa* biofilms. DGL13K displayed in vitro activity against ESBL and KPC-producing *Klebsiella pneumoniae* (MICs 16-32 μg/ml), MDR and XDR *P. aeruginosa*, and XDR *Acinetobacter baumannii* carrying metallo-beta-lactamases (MICs 8-32 μg/ml). *P. aeruginosa* showed low inherent resistance to DGL13K and the increased metabolic activity and growth caused by sub-MIC concentrations of GL13K peptides did not result in acquired bacterial resistance. Daily dosing for approximately two weeks did not increase the MIC of DGL13K or cause cross-resistance between LGL13K and DGL13K. These data suggest that DGL13K is a promising candidate antimicrobial peptide for further development.

## 1. Introduction

Antimicrobial peptides (AMPs) have been considered as an alternative to traditional antibiotics and may represent a different therapeutic modality with reduced opportunity for bacterial resistance (Lazar et al., 2018, Mangoni et al., 2016). The possibility of bacterial resistance to AMPs has been extensively debated. On the one hand, it has been proposed that their mode of action at the cell membrane makes resistance unlikely (Hale and Hancock, 2007, Mangoni et al., 2016, Zasloff, 2002) and peptides such as polymyxin B and nisin have been used for decades with no significant resistance (Martínez and Rodríguez, 2005). On the other hand, resistance can be generated under laboratory conditions (Habets and Brockhurst, 2012, Perron et al., 2006) causing concerns that bacteria that become resistant to a therapeutic AMP would also be resistant to endogenous human host-defense peptides (“arming the enemy”) (Bell and Gouyon, 2003, Dobson et al., 2014, Fleitas and Franco, 2016), as shown for pexiganan and HNP-1 (Habets and Brockhurst, 2012). A recent study suggests that AMPs are more likely to show collateral sensitivity rather than cross-resistance to traditional antibiotics (Lazar et al., 2018). In addition, we have recently reported that closely related peptide enantiomers can show significant differences in their interactions with bacterial defense mechanisms (Bechinger and Gorr, 2017, Hirt et al., 2018).

We previously described the design of anti-inflammatory and bacterial agglutinating peptides based on the sequence of the human salivary protein BPIFA2 (Abdolhosseini et al., 2012b, Geetha et al., 2005, Gorr et al., 2011, Gorr et al., 2008). A modified peptide, GL13K, was developed by substituting three polar or charged amino acids with lysine residues (Abdolhosseini et al., 2012a). The resulting peptide is a more cationic and highly bactericidal peptide, which retains anti-inflammatory activity in vitro and in vivo (Abdolhosseini et al., 2012a). A second generation, D-enantiomer of GL13K resists bacterial proteases (Hirt and Gorr, 2013, Hirt et al., 2018) and is bactericidal against gram-negative and gram-positive bacteria, including vancomycin-resistant *Enterococcus faecalis* and methicillin-resistant *Staphylococcus aureus* (Gorr et al., 2019, Hirt et al., 2018). Interestingly, a similar D-enantiomer selectivity of gram-positive bacteria was reported for the AMP M33-D (Falciani et al., 2012). The goal of this study was to determine the relative activity of the L-and D-enantiomers of GL13K to wild-type and drug-resistant strains of gram-negative bacteria and bacterial biofilms. In addition, we show that *Pseudomonas aeruginosa* exhibit hormesis in response to subinhibitory concentrations of the GL13K peptides but this does not result in acquired resistance to DGL13K or cross-resistance between the L- and D-enantiomers of GL13K.

## 2. Materials and Methods

### 2.1 Bacterial isolate collection

The laboratory strains, *P. aeruginosa* Xen41, a bioluminescent derivate of PA01 (Xenogen, Alameda, CA; now Perkin-Elmer, Waltham, MA), *P. aeruginosa* ATCC 27853, and *K. pneumoniae* ATCC 13883 were used as quality control strains and run in parallel with each MIC experiment. Four clinical *P. aeruginosa* strains (55, 147, 220, 237) collected from Boston, MA, and 2 clinical strains (507, 508) from Philadelphia, PA were tested (Hirsch et al., 2015). Six clinical isolates of *K. pneumoniae* were tested including three from Boston, MA (19, 127, 132) and three from Philadelphia, PA (556, 584, 596). Finally, six *A. baumannii* isolates acquired from the Gram Negative Carbapenemase Detection and *Acinetobacter baumannii* panels of the CDC & FDA Antibiotic Resistance Isolate Bank (Atlanta, GA) (wwwn.cdc.gov/arisolatebank) were tested: AR Bank #33, #52, #102, #280, #290, and #294. The resistance phenotype for each strain is listed in **Table 1**. Isolates were characterized as multidrug-resistant (MDR) if non-susceptible to ≥1 agent in ≥ 3 antimicrobial categories, and extensively-drug resistant (XDR) if non-susceptible to ≥ 1 agent in ≥ 6 antimicrobial categories (Magiorakos et al., 2012).

**Table 1:** LGL13K and DGL13K MIC values determined against wild-type and drug-resistant strains of gram-negative bacteria.

| Species | Strain | Clinical isolate reference | Resistance phenotype | Resistance mechanisms | MIC: LGL13K (µg/ml) | MIC: DGL13K (µg/ml) |
| --- | --- | --- | --- | --- | --- | --- |
| *P. aeruginosa* | PA01 | N/A | Reference |  | 128 | 32 |
|  | ATCC27853 | N/A | Reference ATCC |  | 128 | 32 |
|  | 55 | (Hirsch et al., 2015) | XDR phenotype | ND | 128 | 64 |
|  | 147 | (Hirsch et al., 2015) | MDR phenotype | ND | 128 | 64 |
|  | 220 | (Hirsch et al., 2015) | Wild-type | ND | 128 | 64 |
|  | 237 | (Hirsch et al., 2015) | MDR phenotype | ND | 64 | 32 |
|  | 507 | Present study | MDR phenotype | ND | >128 | 128 |
|  | 508 | Present study | MDR phenotype | ND | 128 | 64 |
| *A. baumannii* | AR Bank #33 | N/A | XDR phenotype | NDM-1, OXA-94, sul2 | 64 | 16 |
|  | AR Bank<br>#52 | N/A | XDR<br>phenotype | OXA-100,<br>OXA-58,<br>sul2 | 64 | 8 |
|  | AR Bank<br>#102 | N/A | XDR<br>phenotype | ADC-25,<br>armA,<br>catB8,<br>mph(E),<br>msr(E),<br>OXA-66,<br>strA, strB,<br>sul1 | 128 | 32 |
|  | AR Bank<br>#280 | N/A | XDR<br>phenotype | aac(3)-Ia,<br>ADC-25,<br>aph(3')-Ic,<br>OXA-66,<br>strA, strB,<br>sul1, TEM-1D | 64 | 32 |
|  | AR Bank<br>#290 | N/A | XDR<br>phenotype | ADC-25,<br>aph(3')-Ic,<br>aph(3')-VIa,<br>armA, catB8,<br>mph(E),<br>msr(E),<br>OXA-23,<br>OXA-66,<br>strA, strB,<br>sul1, TEM-1D | 64 | 32 |
|  | AR Bank<br>#294 | N/A | XDR<br>phenotype | aac(3)-IIa,<br>aph(3')-VIa,<br>OXA-23,<br>OXA-65,<br>strA, strB,<br>sul2, TEM-1B | 64 | 32 |
| *K. pneumoniae* | ATCC<br>13883 | N/A | Reference<br>ATCC |  | 64 | 8 |
|  | 19 | (Hirsch et<br>al., 2015) | Wild-type | ND | 64 | 16 |
|  | 127 | (Hirsch et<br>al., 2015) | Wild-type | ND | 64 | 16 |
|  | 132 | (Hirsch et<br>al., 2015) | ESBL<br>phenotype | ND | 64 | 32 |
|  | 556 | (Chen et al., 2017) | Carbapenem-resistant MDR phenotype | KPC-2 | 64 | 16 |
|  | 584 | (Chen et al., 2017) | Carbapenem-resistant MDR phenotype | KPC-3 | 64 | 16 |
|  | 596 | Present study | ESBL phenotype | ND | 64 | 16 |

ESBL: extended-spectrum beta-lactamase; N/A: not applicable; ND: not determined

### 2.2 Peptides

Polymyxin B was purchased from MilliporeSigma (St. Louis, MO). LGL13K (GKIIKLKASLKLL-NH2) (17) and an all-D-amino acid version of this peptide (DGL13K) (11, 18) were purchased from Bachem AG (Bubendorf, Switzerland). The non-bactericidal control peptide GL13NH2 (GQIINLKASLDLL-NH2) (15, 17) was purchased from Aapptec (Louisville, KY). GL13K peptides were synthesized by Fmoc chemistry and the TCA form isolated at >95% purity by reverse-phase HPLC. The purity and identity of each peptide were verified by the suppliers by reverse-phase HPLC and mass spectrometry, respectively. The lyophilized peptides were re-suspended in 0.01% sterile acetic acid at 10 mg/ml and stored at 4°C. All peptide batches were validated by minimal inhibitory concentration (MIC) testing prior to use, using the modified Hancock protocol described below.

### 2.3 MIC determinations

MICs were determined via two different methods: the broth microdilution (BMD) reference method used for traditional antibiotic susceptibility testing as recommended by the Clinical and Laboratory Standards Institute (28), and the Modified Hancock protocol, a BMD method for cationic AMPs (29).

#### 2.3.1 CLSI BMD protocol

MICs were determined in at least duplicate on separate days. *P. aeruginosa* ATCC 27853 was used as a control strain and run in parallel with each experiment. Briefly, all test isolates and ATCC reference strains were subcultured twice consecutively onto blood agar plates from storage at −80°C and incubated overnight at 35°C. Single isolated colonies were used to inoculate cation-adjusted Mueller-Hinton broth (BBL, Becton Dickinson and Company, Sparks, MD) to a final density of approximately 5 × 10^5^ CFU/ml in each well of a 96-well plate. Bacterial inocula were verified via enumeration following plating of ten-fold dilutions of the inoculum suspension.

#### 2.3.2 Modified Hancock protocol

Broth microdilution assay for cationic antimicrobial peptides (29) was performed as previously described (19). Briefly, a 20 μl solution (1 mg/ml) of each peptide was serially diluted 2-fold in a 1:10 dilution of phosphate-buffered saline (PBS) (Hyclone; GE Healthcare, Pittsburgh, PA) in dH2O (10%PBS), and then mixed with 100 μl of *P. aeruginosa* Xen 41 (10^5^ CFU/ml) in Mueller-Hinton Broth. Final volume in each well was 120 μl and the peptide concentration range tested was 167 μg/ml – 0 μg/ml. Samples were incubated in polypropylene plates at 37°C overnight with gentle shaking. The optical density at 630 nm (OD630) and luminescence were read in a Synergy HT plate reader (BioTek, Winooski, VT) and plotted against peptide concentration. The MIC was read as the lowest peptide concentration that prevented bacterial growth.

### 2.4 Biofilm assay

*P. aeruginosa* Xen 41 (5 x 10^5^ CFU/well, 100 μl LB) were incubated with shaking overnight in 96-well microtiter plates at 37°C. The wells were aspirated and the attached biofilms washed with 200 μl PBS. To each well was added 150 μl MHB or PBS containing a 2-fold serial dilution of peptide (concentration range 1 mg/ml – 1.95 μg/ml). The plates were incubated 60 min at 37°C and luminescence determined in a BioTek plate reader to quantify live cells.

To determine total cells (live+dead) in the attached biofilm, the wells were aspirated and washed with 2 x 200 μl PBS. The plates were incubated with 150 μl/well of 0.03% crystal violet for 30 min at room temperature. The wells were aspirated and washed with 2 x 300 μl PBS followed by 2 x 300 μl dH2O. To each well was added 200 μl 95% ethanol, incubated for 30 min at 37°C, and the OD630 determined. The readings for each peptide were normalized by dividing with the luminescence or OD of the samples with the lowest peptide concentration.

### 2.5 Hormesis

To determine the effect of subinhibitory concentrations of GL13K peptides on bacterial growth and metabolic activity, MIC values (modified Hancock protocol) were read spectrophotometrically and the OD630 (growth) and luminescence (metabolic activity) were determined at each peptide concentration. The peptide concentrations were converted to fold-MIC for each peptide and plotted to allow direct comparison of peptides with different MICs.

### 2.6 Frequency of resistance

LE agarose (BioExpress, Kaysville, UT) was dissolved at 1% in Mueller-Hinton Broth at 95°C. The agarose broth was cooled to 60°C, 100 μg/ml DGL13K was added and the DGL13K-agarose poured in 10 cm petri dishes. Overnight cultures of *P. aeruginosa* Xen41 were pelleted and suspended in sterile 0.9% saline at 5 x 10^8^ CFU/ml (an aliquot was diluted and cultured on agar to validate the concentration of the culture). One ml bacterial culture was plated on each of duplicate DGL13K-agarose plates and incubated overnight at 37°C. Surviving colonies were enumerated as a fraction of 10^9^ plated CFU.

### 2.7 Serial MIC assay

This assay was performed to determine potential development of resistance, as described previously (11). Briefly, an initial MIC assay was prepared using the modified Hancock protocol. The MIC was recorded the following day and the bacteria in the wells containing 0.5xMIC of each peptide (i.e. the highest peptide concentration that allowed growth) were diluted 1000-fold in Mueller-Hinton Broth and 100 μl/well used to inoculate a new MIC plate. The MIC assay was repeated daily for 16 days. On day 15, bacteria that had been exposed to LGL13K were treated with DGL13K to determine cross-resistance.

## 3. Results

### 3.1 Activity against drug-resistant bacteria

Drug-resistant bacteria are an increasing problem and novel antibiotics are urgently needed. The second generation AMP DGL13K has been found to be highly effective against vancomycin-resistant *E. faecalis* and methicillin-resistant *S. aureus* (Gorr et al., 2019, Hirt et al., 2018). In this study, we tested the L- and D-enantiomers of GL13K against drug-resistant strains of the gram-negative bacteria *Klebsiella pneumoniae*, *Acinetobacter baumannii* and *P. aeruginosa* (Table 1).

The MICs for DGL13K are generally lower than those recorded for LGL13K, in agreement with our previous results (Gorr et al., 2019, Hirt and Gorr, 2013). Against the three gram-negative species tested, DGL13K was most active against *K. pneumoniae* and *A. baumannii*, with MICs ranging from 8-32 μg/ml (Table 1)._Against *K. pneumoniae* isolates, MICs for ESBL- and KPC-producing strains did not significantly differ when compared to the ATCC reference strain. Against *P. aeruginosa*, MICs were within 2 doubling dilutions for MDR and XDR isolates (32-128 μg/ml) when compared to the reference strains (32 μg/ml). The latter results are about 6-fold higher than those previously reported for *P. aeruginosa* (Gorr et al., 2019)

### 3.2 Activity against biofilms of *P. aeruginosa*

LGL13K and DGL13K kill biofilms of *P. aeruginosa* (Hirt and Gorr, 2013). To compare the dose needed to kill biofilms with the MIC, biofilms were incubated with increasing doses of LGL13K, DGL13K, GL13NH2 and the control antimicrobial peptide polymyxin B. **Fig. 1A** shows that 99% reduced viability (LD99) of wild-type *P. aeruginosa* was achieved at a concentration of 32 μg/ml, whereas 128 μg/ml of LGL13K or polymyxin B were required to reach a comparable reduction of viability. Thus, the LD99 for biofilms is similar to the MIC achieved for both GL13K enantiomers (**Table 1**). Biofilm viability was not affected by the control peptide GL13NH2, which is not bactericidal (Abdolhosseini et al., 2012a). The biomass of the biofilms was not reduced by the peptide treatments (**Fig. 1B**), suggesting that the dead bacteria remained attached to the substrate under these conditions.

**Figure 1:**
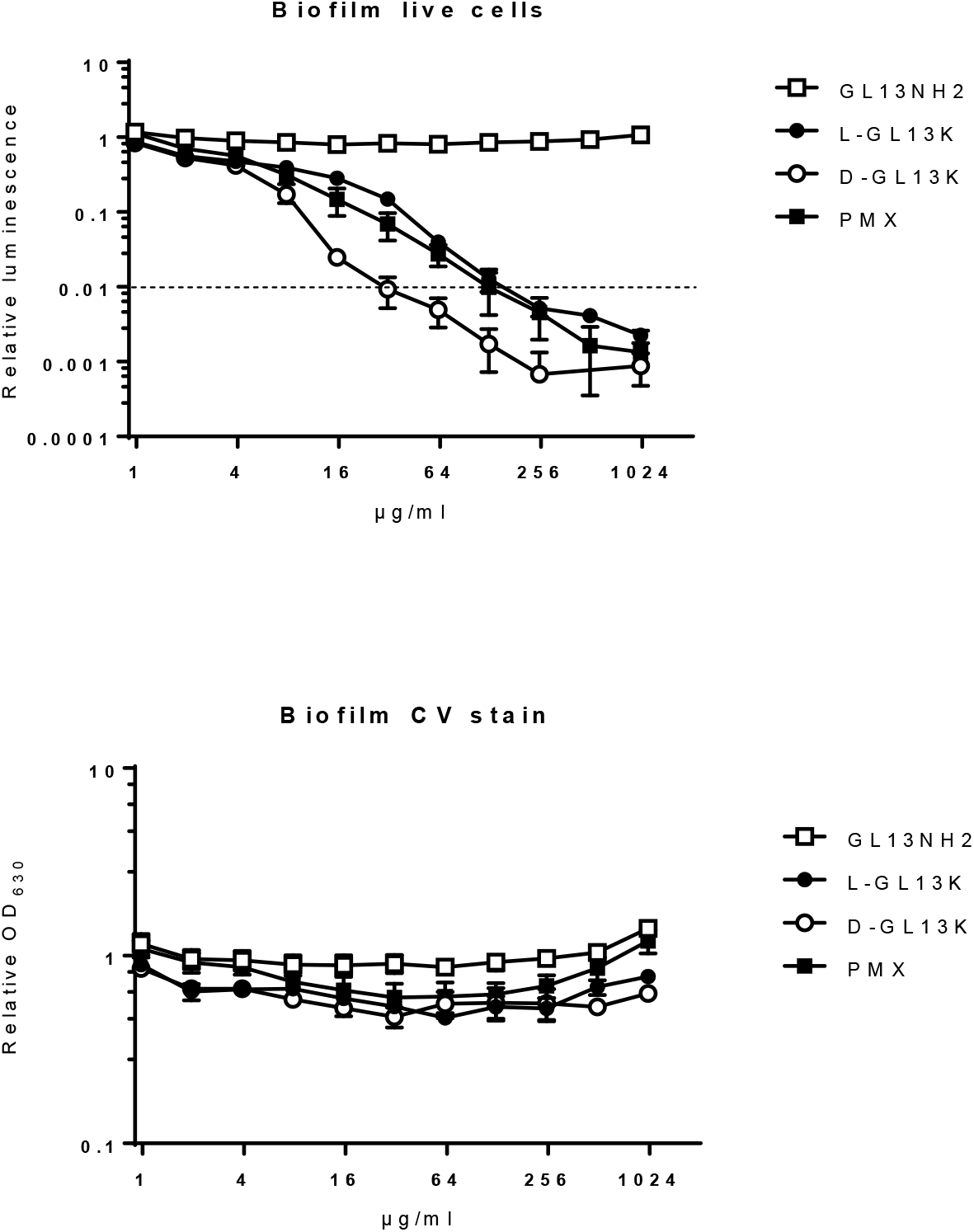
Bactericidal activity of peptides against *P. aeruginosa* biofilms. Biofilms were incubated for 1h with peptides, at the concentrations shown. **A.** live cells were quantitated by luminescence. Dotted line indicates 99% killing of biofilm. **B.** Biofilm biomass was quantitated by crystal violet staining. PMX = polymyxin B. Data from two independent experiments performed in duplicate were normalized to the mean of the lowest peptide concentration in each experiment. Data shown as mean ± SEM (N=4).

### 3.3 Effect of sub-inhibitory peptide concentrations on bacterial growth

In dose response experiments with *P. aeruginosa*, we noted that the OD_630_ of the cultures increased with increasing peptide concentration, up to 0.5 x MIC. A similar effect of subinhibitory concentrations has been reported for many toxic substances (hormesis) (Stebbing, 1982), including traditional antibiotics (Wang et al., 2017). To further evaluate this effect, the growth of *P. aeruginosa* was determined by culture density (OD_630_) and bacterial metabolic activity was determined by cellular luminescence (Robinson et al., 2011). Increasing peptide concentration up to 0.5 x MIC increased the OD_630_ by about 50% (**Fig. 2A**) while metabolic activity increased 2-4-fold (**Fig. 2B**).

**Figure 2:**
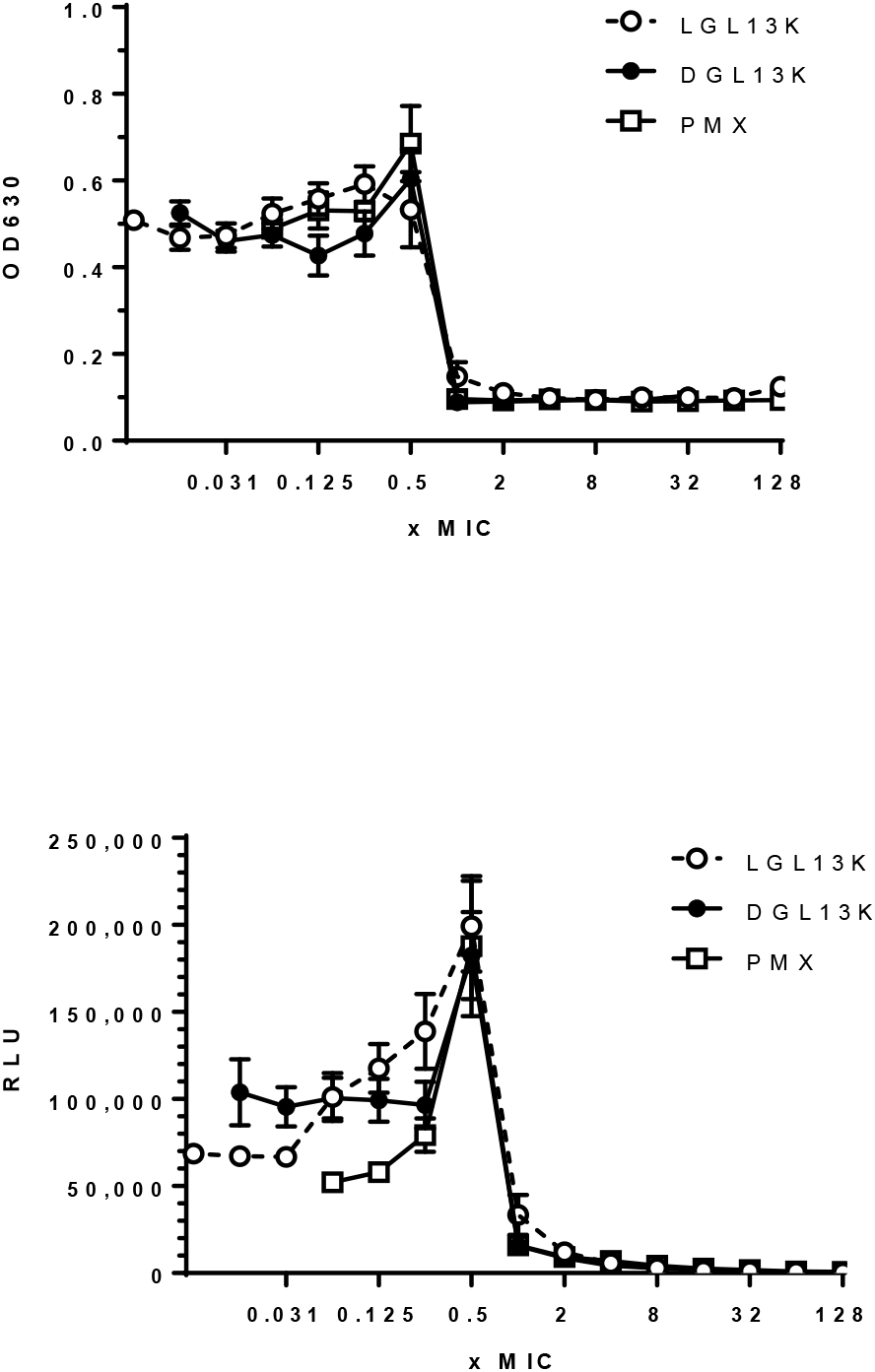
*P. aeruginosa* culture response to increasing peptide concentrations. *P. aeruginosa* Xen 41 were incubated with increasing concentrations of LGL13K (open circles), DGL13K (closed circles) or polymyxin B (PMX, open squares). The data from 2-3 independent experiments are shown as mean ± SEM, N=4-8.

### 3.4 Bacterial resistance to GL13K

The frequency of resistance of *P. aeruginosa* was less than 10^-9^ when the bacteria were plated on agar containing 100 μg/ml DGL13K, i.e. 3xMIC determined in **Table 1**. Thus, these bacteria show very low inherent resistance to DGL13K.

The increased metabolic activity caused by culturing *P. aeruginosa* in the presence of 0.5xMIC of the GL13K peptides raised the question if the bacteria acquire resistance to the GL13K enantiomers when they are cultured under sub-inhibitory peptide concentrations. Repeated exposure of *P. aeruginosa* to 0.5xMIC of DGL13K did not increase the MIC of this peptide after 16 rounds (days) of selection (**Fig. 3**). The MIC for LGL13K trended towards a 2-fold increase but this did not reach statistical significance (P<0.06). Importantly, bacteria that had reached the higher MIC for LGL13K did not show an increased MIC for DGL13K (**Fig. 3**, closed square). The lack of cross-resistance between these two closely related AMP enantiomers is promising for future clinical use (Fleitas and Franco, 2016).

**Figure 3:**
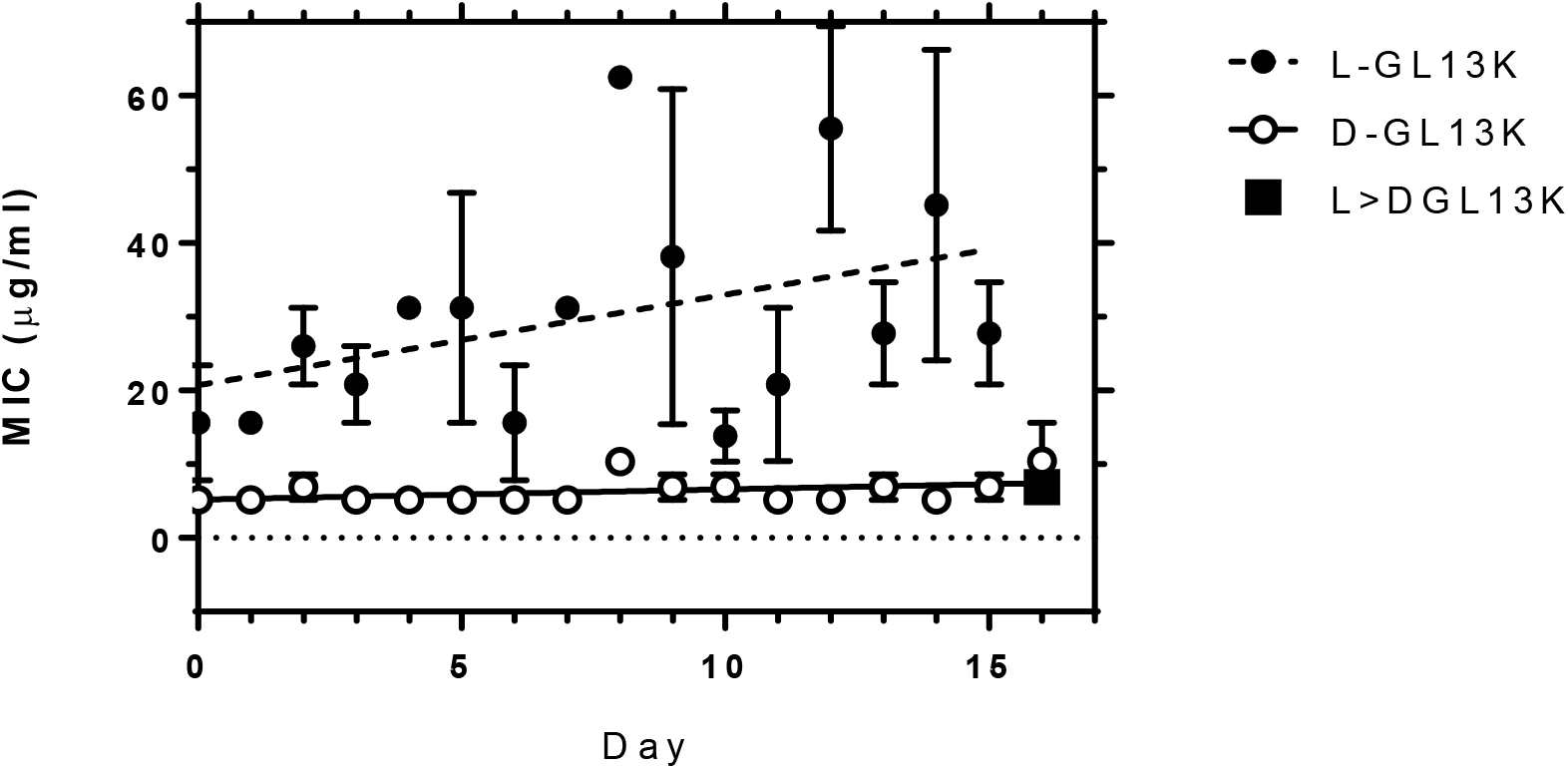
**Development of resistance in *P. aeruginosa*** by repeated treatment with LGL13K (closed circles) or DGL13K (open circles). The MIC determinations were plotted for each day (mean ± SEM, N=3) and analyzed by linear regression. Samples treated with LGL13K for 15 days were then treated with DGL13K and the MIC determined (square. Mean ± SEM, N=3).

## 4. Discussion

The antimicrobial peptide enantiomers LGL13K and DGL13K have shown promising activity against gram-negative (LGL13K and DGL13K) (Abdolhosseini et al., 2012a, Hirt and Gorr, 2013) and gram-positive bacteria (DGL13K) (Hirt et al., 2018). In this report, we determined the antibacterial activity against additional bacterial species as well as several drug-resistant strains of clinically important gram-negative bacteria. The second-generation antimicrobial peptide DGL13K shows activity against the tested drug-resistant strains that is similar to that of the corresponding wild-type strains. Importantly, DGL13K displayed activity against isolates of multiple species of resistant gram-negative pathogens including ESBL and KPC-producing *Klebsiella pneumoniae* (MICs 16-32 μg/ml), MDR and XDR *P. aeruginosa*, and XDR *A. baumannii* carrying metallo-beta-lactamases (MICs 8-32 μg/ml). Treatment options for infections caused by these resistant pathogens are limited and often result in treatment with more toxic agents such as the polymyxins or aminoglycosides (Butler et al., 2019, Hirsch and Tam, 2010). DGL13K did not exhibit collateral sensitivity (Lazar et al., 2018) in drug-resistant strains of gram-negative bacteria. Similarly, we have recently reported that the MIC for drug-resistant *Staphylococcus aureus* and *Enterococcus faecalis* are not lower than the MIC for wild-type strains. Indeed, bactericidal activity is similar to the growth inhibiting activity in most cases and the peptide is highly active against established bacterial biofilms. Thus, DGL13K is a promising candidate for further development.

The data in Table 1 were generated using the CLSI protocol for broth microdilution (Clinical and Laboratory Standards Institute, 2019) while our previous results (Abdolhosseini et al., 2012a, Gorr et al., 2019, Hirt and Gorr, 2013) were obtained with a modified version of the protocol developed for cationic antimicrobial peptides by Hancock (Hancock, 1999). Similarly, it has been reported that the ‘Hancock protocol’ results in lower MICs for cationic peptides than the protocol described by CLSI (Giacometti et al., 2000). In direct comparisons, the MIC of DGL13K was not affected by plate materials (polystyrene vs. polypropylene), incubation with or without shaking or inoculum 10^5^ CFU/ml vs. 10^6^ CFU/ml) (data not shown). Thus, the exact cause of the higher MIC values for the CLSI protocol is not clear but likely results from a combination of factors.

In the course of antibacterial activity studies, we noted that subinhibitory (sub-MIC) concentrations of the antimicrobial peptides caused increased growth and metabolic activity of *P. aeruginosa*, which was most notable at 0.5xMIC. A similar phenomenon (hormesis) has been described for several toxins in multiple species and taxa (Mattson, 2008, Stebbing, 1982). Interestingly, the effect in bacteria has been linked to “medium-richness”. Thus, the growth promoting effect of the antibiotics sulfamethazine and erythromycin were more pronounced in dilute MHB than in full strength MHB and the effect was absent in LB medium (Calabrese et al., 2010). The effect of GL13K peptides was stronger on metabolic activity than overall growth, suggesting that *P. aeruginosa* were selectively stimulated in metabolic pathways, although the mechanism of stimulation is not yet known.

Re-analysis of peptide dose-response curves (MIC assays) for the gram-positive bacteria *Streptococcus gordonii, Enterococcus faecalis* (Hirt et al., 2018) and *Staphylococcus aureus* (Gorr et al., 2019) revealed that some strains exhibited a similar growth stimulatory effect at subinhibitory concentrations of GL13K peptides (Gorr, unpublished). DGL13K, but not LGL13K, circumvents cell wall defense mechanisms that include D-alanylation of teichoic acids. Interestingly, the hormesis effect was observed in D-alanylation mutants that had lost resistance to LGL13K, suggesting that the effect is not associated with the initial point of attack at the cell surface. A better understanding of the cellular targets for GL13K peptides in gram-negative and gram-positive bacteria will be needed to determine the different mechanisms that allow peptide-induced bacterial hormesis.

*P. aeruginosa* show very low inherent resistance to DGL13K and the increased metabolic activity and growth at sub-MIC concentrations of GL13K peptides did not result in acquired bacterial resistance. Daily dosing for about two weeks had no effect on the MIC of DGL13K towards *P. aeruginosa*. Similar results were recently reported for the gram-positive bacteria *S. gordonii* and *E. faecalis* treated with DGL13K (11). Together, these results support the notion that, although bacteria can use surface modifications to block the attack by antimicrobial peptides, the killing ultimately depends on membrane disruption (12), which may be costly to combat through bacterial mutation (3). It has been proposed that host antimicrobial peptides have remained effective against invading pathogen through a process of co-evolution (33). The lack of inherent or acquired resistance of gram-negative (this study) and gram-positive bacteria (11) and the ability of DGL13K to overcome bacterial defense mechanisms that affect LGL13K give hope that antimicrobial peptides can be designed to address bacterial resistance without “arming the enemy”.

## 5. Acknowledgement

SUG gratefully acknowledges research funds provided by the University of Minnesota School of Dentistry.

